# Wing pitch timing and wing elevation modulate forces and body pitch in forward flapping flight

**DOI:** 10.64898/2026.03.06.710029

**Authors:** Victor Colognesi, L. Christoffer Johansson

## Abstract

Aerodynamic performance in flying animals can be controlled not only by changes in wing shape and size, but also by changes in wing motion. In flapping wings, the two wings interact aerodynamically, where the interaction depends on wingbeat phase, wing proximity, and wing attitude. Although, observations of free-flying animals show wings are relatively close during forward flight, how these interactions influence force production remain unclear. Here we used a robotic flapping wing and quantitative flow measurements to test the effect of wing interactions while varying mean wing elevation and pitch timing during upstroke–downstroke transitions. We show that both force magnitude and direction depend strongly on these parameters and their interaction. The fitted response models showed that high wing position combined with early pitching maximized vertical force, whereas low-to-mid wing position with synchronized-to-late pitching increased thrust. Efficiency metrics were highest near mid-to-high wing position combined with late pitching. In addition, we found that transition phases strongly affected thrust generation and produced substantial body pitch torques. These findings demonstrate that small kinematic adjustments can markedly alter aerodynamic performance and be used for tailless flight control. This offers mechanistic explanations for observed animal wing motions and novel strategies for controlling flapping drones.

## Introduction

Animals in flapping flight regulate both the magnitude and direction of aerodynamic forces through passive wing deformations and active muscular control. Birds, bats, and insects continually adjust wing trajectory, attitude, and velocity to maintain stability (1), modulate speed (2–8), and execute maneuvers (9–12). Variations in kinematics can disproportionately influence both the magnitude and orientation of aerodynamic forces (13,14), thereby shaping flight performance across speeds, accelerations, and environmental conditions. This flexibility arises from the biological flight apparatus, which allows for virtually infinite kinematic variation. Understanding how specific wing-motion parameters affect force generation is therefore critical both for explaining the diversity of natural flight behaviors and for guiding the design of the next generation flapping-wing robots (15).

Despite major advances in animal-flight research (16–20), the aerodynamic consequences of most kinematic variation remain unresolved. The link between kinematics and aerodynamics is relatively well understood for hovering flight (14,21–24), while prior work on forward-flight remains more limited. Work on forward flapping flight has primarily examined parameters such as Strouhal number (flapping velocity relative to forward speed) (25) and wing pitch amplitude (26), or simplified flapping to heaving wings (27,28), which is insufficient to capture the 3D flow phenomena of flapping wings (29). However, studies of hovering flight dynamics and theoretical reasoning suggest additional strategies to control aerodynamics, which may apply to forward flight but remain largely unexplored. Indeed, recent experiments showed that mean wing elevation, a previously untested parameter, can modulate both force magnitude and orientation by altering wing-wing interactions during the downstroke (30). Another particularly potent parameter to control forces is pitch timing, which influences the angle of attack and wing-wake interactions in hovering-flight studies (14,31,32). In forward-flying animals, pitch timing and mean wing elevation have rarely been explicitly studied (but see (33) for a 2D study), although available evidence suggest that the parameters vary with flight speed in flying animals (4,8,34–39). However, the mechanistic links between these kinematic parameters and the resulting aerodynamic forces are difficult to isolate in vivo. A systematic parametric analysis is therefore needed to disentangle how variation in wing kinematic parameters, individually and in combination, affects lift, thrust, force orientation, efficiency, and pitch-control in forward flapping flight.

Here, we address this knowledge gap by experimentally quantifying how wing pitch timing and mean wing elevation interact to shape aerodynamic performance using a robotic flapping wing. Using a wind tunnel setup with particle image velocimetry (PIV) to measure airflow, we independently manipulate wing pitch timing and mean wing elevation and quantify the resulting lift, thrust, force orientation, and energetic costs. This approach isolates how these parameters alter aerodynamic output, offering a mechanistic framework for interpreting kinematic variability in flying animals. By revealing how targeted timing and positional adjustments regulate aerodynamic forces, our findings offer new insights into natural flight control and inform strategies for achieving robust, responsive, and energy efficient control in flapping drones.

## Material and methods

### Experimental setup

We used a robot (previously described in (30,40)), fitted with a novel 3D-printed wing (wing length = 150 mm, mean chord = 39 mm, maximum thickness = 6 mm, see Fig. S1). The robot is capable of independently controlling the angular elevation of the wing above (max 90 degrees) and angular depression of the wing below (max 90 degrees) the horizontal plane through the flapping axis, the wingbeat frequency (*f*, in the range 0-15 Hz) and the pitch angle (γ, varying ±40 degrees). These factors can be controlled as continuous functions over time and essentially take any shape allowed by motor acceleration. Here we controlled the mean wing elevation (*MWE*) and the pitch angle (γ). *MWE* was defined as the average of the maximum elevation and depression angles of the wing relative to the horizontal plane. Positive values represent wingbeats centered above the horizontal, negative below. Pitch angle was defined as the angular rotation of the wing around the spanwise axis. The robot was mounted on the wall of the wind tunnel and was controlled using a custom software via a microcontroller (30,40).

For the experiments we used fixed tunnel speed (7.2 m/s), tip-to-tip angular amplitude of the flapping (*A*_ang_ = 70 degrees), flapping frequency (6.7 Hz) and downstroke ratio (0.5, equal duration of down and upstroke). For all experiments, the wing flaps at a constant velocity and constant pitch for the downstrokes (-4 degrees), and upstrokes (14 degrees). At the transitions, both the velocity and the pitch follow a sinusoidal trajectory, taking 10% of the total flapping period, ensuring a continuous and smooth transition. The timing of the pitching (*TP*) was then shifted relative to the turning points with a delay of minus half of the transition time (so -5% of the period), 0 and plus half of the transition time (Fig. S2). An ‘early’ pitch means the wing rotates before reaching the turning point, while a ‘late’ pitch means rotation occurs after the turning point. We varied the *MWE* (by independently controlling the maximum elevation and depression angle of the wing) between -30 deg (low) and 30 deg (high) in steps of 30 degrees. For each kinematic condition, we made three measurements, where each measurement included four flapping cycles. The wing motion was stopped between each measurement and subsequently started again providing unique starting conditions for the wing for each sequence. We therefore considered the measurements to be statistically independent.

We defined a right-handed coordinate system with *x* in the tunnel flow direction and *z* in the upwards direction. We measured the flow using stereo particle image velocimetry (sPIV) in a plane perpendicular to the freestream flow (*yz*-plane) of the Lund University wind tunnel (41). The sPIV system (Lavision, Göttingen, Germany) includes a dual pulse laser (LDY304PIV laser, Litron Lasers, Rugby, England), operated at f_L_=720 Hz, four high speed cameras (Photron Nova R2, Photron Deutschland GmbH, Reutlingen, Germany) and acquisition and analysis software (Davis 10.2.1, Lavision, Göttingen, Germany). The cameras were set up to all view the same area (∼340 x 400 mm, *yz*). The analysis settings were; pre-processing with a subtract minimum over time (5 frames), followed by a vector calculation with an initial box size of 96×96 and a final box size of 24×24, with 50% overlap. The final vector field was subjected to a vector validation of strength 2. To remove erroneous vectors and noise we did a final post processing step. This included a 1x universal outlier detection, removing vectors deviating more than 2.5 standard deviation from their neighbors (7×7 box) and reinserting the second-best vector if less than 3 standard deviations from their neighbors. This step also included interpolating missing vectors and a 5×5 denoising routine. The resulting vector field was 150×177 vectors with a spacing of 2.27 mm.

### Analysis

The vector fields were analyzed using custom written software in Matlab R2022a (Mathworks Inc.). The data were masked to remove erroneous vectors at the edge of the vector field and mirrored in the sagittal plane of the robot, to generate a wake representing both wings of an animal. Mirroring the vector field is motivated by the wind tunnel wall acting as an aerodynamic mirror (42–45), making it possible to study wing-wing interactions using a single wing (44). This procedure ignores effects of the boundary layer but given that the boundary layer in the current tunnel is thin (46) and that all measurements are performed at the same speed we consider these effects to be small. In each PIV sequence, the free-stream velocity *U*_∞_ was estimated by averaging the measured streamwise velocity in an area unaffected by the wake of the wing. We then stacked the vector fields in the streamwise direction with a spacing of *U*_∞_/*f*_L_ and estimated the vorticity (*ω*) as the curl of the flow velocities in the volume.

The vertical force, lift (*L*), was then calculated according to,

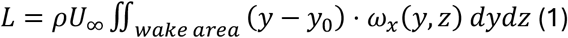

where ρ is the air density, y is the spanwise position, y_0_ the spanwise position of the center of the body and ω_x_ is the vorticity along the freestream axis.

We estimated the thrust (*T*) in the direction of flight as,

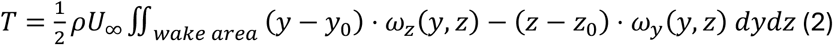

where *ω*_z_ is the vorticity along the vertical axis, *ω*_y_ is the vorticity along the spanwise axis, z the vertical position and *z*_0_ the vertical position of the center of the body (47).

For the comparisons in this paper, we used the time-averaged force over four wingbeats. The direction of the resultant force was determined as the average thrust to average lift ratio (*T*/*L*) and the ability to generate force was determined by estimating the force coefficients,

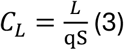

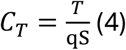

where q is the dynamic pressure 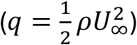 and *S* is the wing area.

We also estimated the useful force (*F*_use_) as the magnitude of the vector sum of *T* and *L* and normalized according to,

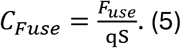

We determined the kinetic energy in the wake as a measure of the cost of generating the forces (48,49), using the following equation:

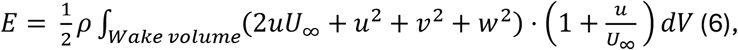

where *u, v, w* are the induced velocities in *x, y, z*, respectively. Contrarily to the vorticity field, which is compact, the velocity field is not captured in its entirety by the PIV measurements. The first term in the energy equation is related to the momentum change due to non-zero thrust production by the wing. While it vanishes for flying objects at force equilibrium this term is necessary to account for the net forces generated by the wing (50). To improve our estimation of the kinetic energy, we performed a Helmholtz-Hedge decomposition and extended the velocity field beyond the dimensions of the wind tunnel (here to a size of 1.2 x 1.2 m cross section) (48,49) prior to estimating the kinetic energy. The reconstruction is based on a masked vorticity field, with the mask generated automatically from the data. The masking started with the large vortical structures, then included a set volume around them. The final mask applied to the 3D vorticity field was an extrusion of a 2D mask normal to the flow direction obtained by combining all 2D slices of the initial 3D mask. Due to the flow field being reconstructed from the vorticity based on the stacked 2D-slices of the flow, velocity at the edges of the volume did not always become zero. We therefore subtracted the mean streamwise velocity at the edges of the volume before calculating the kinetic energy. Assuming no pressure work on the Treftz plane, the power was subsequently estimated by dividing the kinetic energy in the wake by the number of wingbeats (N_wb_= 4) and multiplying by the wingbeat frequency (*f*):

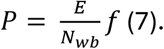

We then normalized power by calculating the power coefficient (C_P_) as,

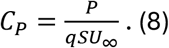

Since the force generation differs between the kinematic settings, we estimated performance parameters to allow for a comparison between the cases. A traditional measure of performance is the cost of transport (COT), a measure of the relative cost of generating *L*, defined as the energy required to transport one unit of weight, one unit of distance. By equating the weight with the lift generated by the wing COT becomes:

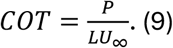

We also determined the power factor (PF)(23,51), which is an efficiency measure of the combined useful force generated. The power factor originates from the actuator disk model and the fact that force is proportional to the induced velocity squared, while power is proportional to the induced velocity cubed. PF was calculated as:

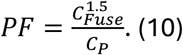

Finally, we computed the pitch torque resulting from the wing thrust. The moment arm used for the torque was computed as follows:

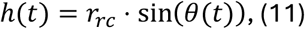

where θ is the flapping angle. We approximated the application point of the thrust along the wing to be located at a distance *r*_*rc*_ from the rotation point of the wing, defined as

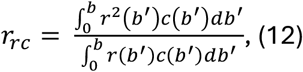

where *b* is the wingspan and *b’* is the spanwise position along the wing. This distance was weighed by the chord, as more force is produced where the wing is wider, and by the radius *r* from the rotation point, accounting for an increase of thrust towards the wing tips.

The flapping angle (*θ*) throughout the wingbeat was determined from the imposed kinematics. To synchronize the wing trajectory with the forces observed in the wake, we evaluated the relative shift within one sequence. We first selected a PIV sequence where the tip vortex was easily identifiable and manually selected the frame where it reached the lowest position. We then determined the frame where the wing was at the end of the downstroke in footage of the wing taken simultaneously to the PIV sampling. The delay between the PIV frame and the corresponding end of downstroke in the wing footage was then applied to every sequence, assuming that the different cases, given the fixed distance between wing and measurement plane and the constant tunnel speed, did not affect the time for the wake to travel from the wing to the PIV plane.

The moment was finally computed as

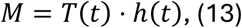

and made dimensionless with

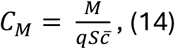

where 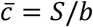 is the mean aerodynamic chord of the wing.

### Statistics

The experimental design sampled a two-dimensional kinematic parameter space with TP and MWE. While limited to three levels per variable, this design allowed us to fit quadratic response surfaces and test how aerodynamic outputs vary across the sampled domain. For this purpose we performed a GLM analysis (*fitglm* function with ‘quadratic’ model, Matlab R2025a, Mathworks Inc., USA), for each of the dependent variables with *TP* and *MWE* as independent, continuous, variables set to values between –1 and 1. Variation is given as standard error. Interaction plots of the fitted quadratic GLM were used to visualize the effect of *TP* across *MWE*.

The GLM results were interpreted as response surfaces over the sampled TP–MWE parameter space, and we quantified if each output was governed primarily by linear trends, curvature, or TP*MWE interaction. For this purpose, we performed hierarchical nested-model comparisons (i.e. incrementally removing one term at a time from the model) and quantified the incremental explanatory power of each block of terms (R^2^_partial_ = (SSE_simple_ - SSE_complex_)/SSE_simple_).

## Results

### Lift

The wingbeat average *C*_L_ varied non-linearly with both *TP* (*TP*: 0.0101±0.0044, p=0.0313; *TP*^2^: - 0.0331±0.0076, p=0.000286) and *MWE* (*MWE*: 0.1466±0.0044, p=33.3 1.14e^-19^; *MWE*^2^: - 0.0664±0.0076, p=2.02e^-08^). There was an increase in *C*_L_ with increasing *MWE*, regardless of *TP* (Fig. 1a). *TP*, per se, had a significant, but weak, effect on *C*_L_. The response surface for *C*_L_ showed a strong interaction effect between *TP* and *MWE* (*MWE*TP*: -0.1071±0.0054, p=4.25e^-15^) (Fig. S3a, g). Increasing *MWE* increased lift across the sampled domain, but the effect of *TP* changed sign with *MWE*: at low *MWE*, earlier *TP* reduced *C*_L_, whereas at high *MWE*, earlier *TP* increased *C*_L_. Thus, *TP* cannot be interpreted independently of *MWE* for *C*_L_ (Figs 1a, S5a).

**Figure 1.**
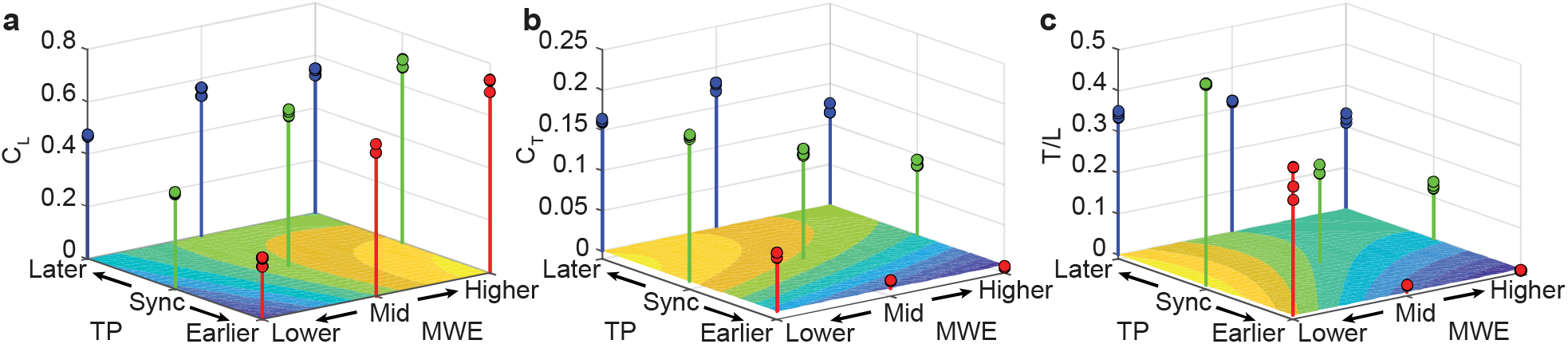
*C*_L_ and *C*_T_ during a wingbeat varies with timing of pitch (*TP*) and mean wing elevation (*MWE*). (a) *C*_L_ increases with increasing *MWE*. The increase is faster when *TP* occurs early relative to the transition between down and upstroke, than when *TP* occurs later relative to the stroke transition. (b) *C*_T_ increases as *MWE* decreases and as *TP* occurs later relative to the transition between down and upstroke. (c) *T*/*L* represents the direction of the force generated by the wing. Highest *T*/*L* is achieved when *MWE* is low and the *TP* is synchronized with the flapping motion, while the lowest *T*/*L* is found when *MWE* is high and *TP* is early. The data points are the wingbeat averages for each measured sequence, and the contour plot shows the output from the glm-model fitted to the data.

The instantaneous *L* over the wingbeat showed during which phases of the wingbeat we found differences between the treatments. In both the *TP* and *MWE* changes, most between-condition differences arise during stroke-reversal phases (Figs 2 and S4). This identifies the turning phases as the main window in which TP and MWE modulate aerodynamic output. Earlier *TP* and higher *MWE* both tended to increase *C*_L_ during this phase. At the transition between upstroke and downstroke we also found the curves diverging, but here the effects of the two kinematic variables differed. Earlier *TP* resulted in more negative lift (reflected in reversed vortex ring during the end of upstroke, Fig. S5), while higher *MWE* resulted in less negative *C*_L_. We also found another effect of *MWE*, where *C*_L_ during the early and mid-downstroke was lower the higher the *MWE*.

**Figure 2.**
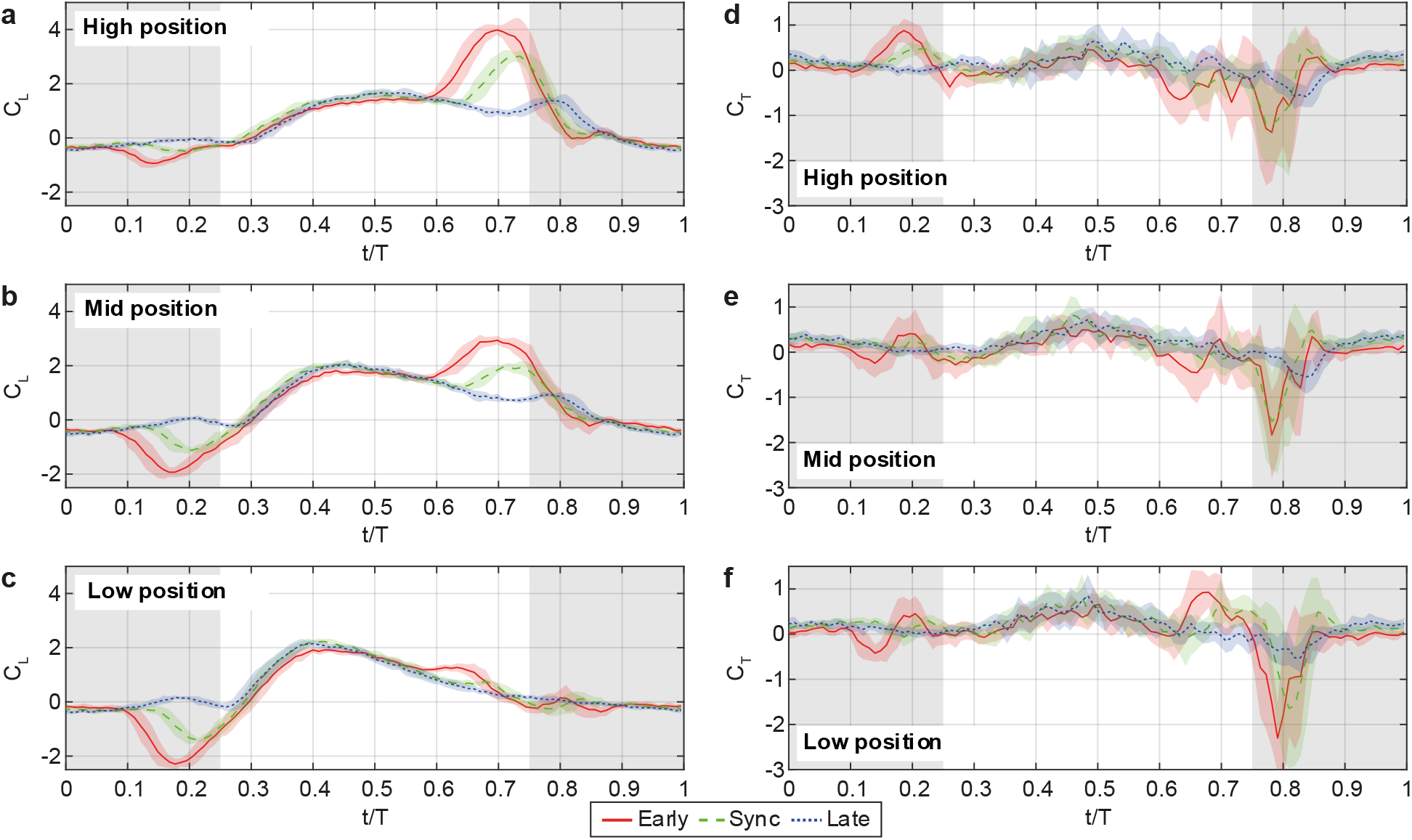
Instantaneous force coefficients through the flapping cycle. The left panels (**a-c**) show the impact on *C*_L_ and the right panels (**d-f**) show the impact on *C*_T_ for the timing of the pitch (*TP*) at different wing positions (*MWE* = High, Mid and Low). In each curve, the line represents the phase-averaged value over the three repeats and four wingbeats of each case, and the shaded area has a width of two standard deviations centered around the mean. Shaded background shows the upstroke.

### Thrust

The wingbeat average *C*_T_ also varied with both *TP* and *MWE*. For *C*_T_, the fitted surface showed strong main effects of both *TP* and *MWE*, with increasing *C*T with decreasing *MWE* (*MWE*: -0.0345±0.0037, p=6.58e^-09^), regardless of *TP* (Fig. 1b). There was also a non-linear effect of *TP* on *C*T (*TP*: 0.0631±0.0037, p=9.22e^-14^; *TP*^2^: -0.0439±0.0064, p=9.14e^-07^), where *C*_T_ increased more the earlier the *TP*. There was no interaction effect between the two factors (p=0.11) (Fig. S5b), indicating that these two kinematic variables contribute additively rather than interactively to thrust production (Fig. 1b).

The instantaneous *C*_T_ over the wingbeat showed that later *TP* reduced negative *C*_T_ during the transition between downstroke and upstroke, but also lower *C*_T_ during the upstroke and downstroke transition (Figs 2 and S4). Lower *MWE* resulted in higher *C*_T_ during the downstroke to upstroke transition.

### Thrust to lift ratio

The force-orientation metric *T*/*L* decreased with increasing *MWE* and increased with later *TP*, with a significant *TP*MWE* interaction reflecting a combination of the trends described above (Fig. 1c). There was a non-linear decrease in *T*/*L* with increasing *MWE* (*MWE*: -0.1342±0.0121, p=3.31e^-10^; *MWE*^2^: 0.0638±0.0210, p=0.0063), regardless of *TP*. There was also a non-linear increase of *T*/*L* with *TP* (*TP*: 0.0915±0.0121, p=2.14e^-7^; *TP*^2^: -0.0789±0.0210, p=0.0012), and a significant interaction effect between the two factors (*MWE*TP*: 0.0496±0.0149, p=0.0032) (Fig. S5c, i). The resulting surface indicates a controllability tradeoff: shifting toward high *MWE* and early *TP* rotates the resultant force upward, whereas shifting toward low *MWE* and synchronized/later *TP* rotates it forward.

### POWER

*C*_P_ varied linearly with *TP* (*TP*: -0.0459±0.0019, p=1.15e^-16^) and non-linearly with *MWE* (*MWE*: 0.0117±0.0019, p=5.12e^-06^; *MWE*^2^: -0.0105±0.0034, p=0.00497) (Fig. 3a), while there was no significant interaction between *MWE* and *TP* (*MWE*TP*: p=0.0628) (Fig. S5d, j). The earlier the pitch occurred the higher the *C*_P_. For *MWE* there is weak increase in *C*_P_ with increasing *MWE*, but with a weak peak in *C*_P_ for the mid wing elevation.

**Figure 3.**
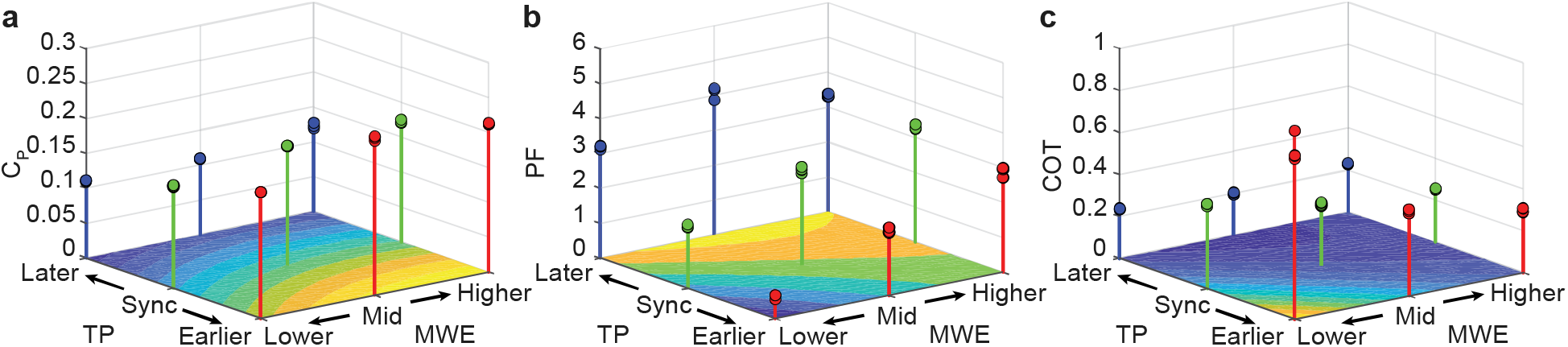
Aerodynamic power and efficiency vary with timing of pitch (*TP*) and mean wing elevation (*MWE*). (a) Power coefficient (*C*_P_) varies strongly with *TP*, increasing the earlier the pitch occurs relative to the transition between downstroke and upstroke. There is also a generally lower *C*_P_, the lower the *MWE*. (b) Power factor (*PF*), a measure of the force/power generated, increases the later the *TP* occurs and the higher the *MWE* (except for the highest *MWE* with the latest *TP*). (c) Cost of transport (*COT*), the energy required to move one Newton of weight one meter, decreases with increasing *MWE* and delayed *TP*. The data points are the wingbeat averages for each measured sequence, and the contour plot shows the output from the glm-model fitted to the data.

### Efficiency

Efficiency metrics exhibited strong *TP*\**MWE* interactions (Fig. S5 d-e, k-l). Late *TP* consistently improved energetic performance, but the benefit of increasing *MWE* depended on *TP*: at early *TP*, higher *MWE* improved *PF* and reduced *COT*, whereas at late *TP* the same change in *MWE* produced smaller gains or even the opposite trend. Thus, efficient force production emerges from a specific combination of kinematic timing and elevation, not from either parameter alone.

For the specific efficiency measures we saw that *PF* varied linearly with *TP* (TP: 0.8681±0.0484, p=3.32e^-14^) and non-linearly with *MWE* (*MWE*: 0.6532±0.0484, p=8.24e^-12^; *MWE*^2^: -0.3729±0.0839, p=0.00022) (Fig. 3b). There was a strong interaction effect (*MWE*\**TP*: -0.5256±0.0593, p=1.54e^-08^) so that when *TP* occurred early *PF* increased strongly with increasing *MWE*, but when *TP* occurred late *PF* decreased with increasing *MWE* (or peaked at mid *MWE*).

*COT* varied non-linearly with *TP* (*TP*: -0.1418±0.0106, p=9.81e^-12^; *TP*^2^: 0.0469±0.0183, p=0.0185) and non-linearly with *MWE* (*MWE*: -0.1118±0.0106, p=7.76e^-10^; *MWE*^2^: 0.0740±0.0184, p=0.000611) (Fig. 3c). There was a strong interaction effect (*MWE*\**TP*: 0.1280±0.0130, p=2.53e^-09^) so that *COT* was highest when *MWE* was low and *TP* occurred early and was lowest when *MWE* was high and *TP* occurred late.

### Body pitch torque

We found the highest values of the pitch torque, *C*_M_, acting on the body when the wing tip was the furthest from the center of the body (Fig. 4). This was the time during the wingbeat that the thrust had the greatest moment arm for pitch generation, but it also coincided with the phase of the wingbeat when we found peaks of thrust.

**Figure 4.**
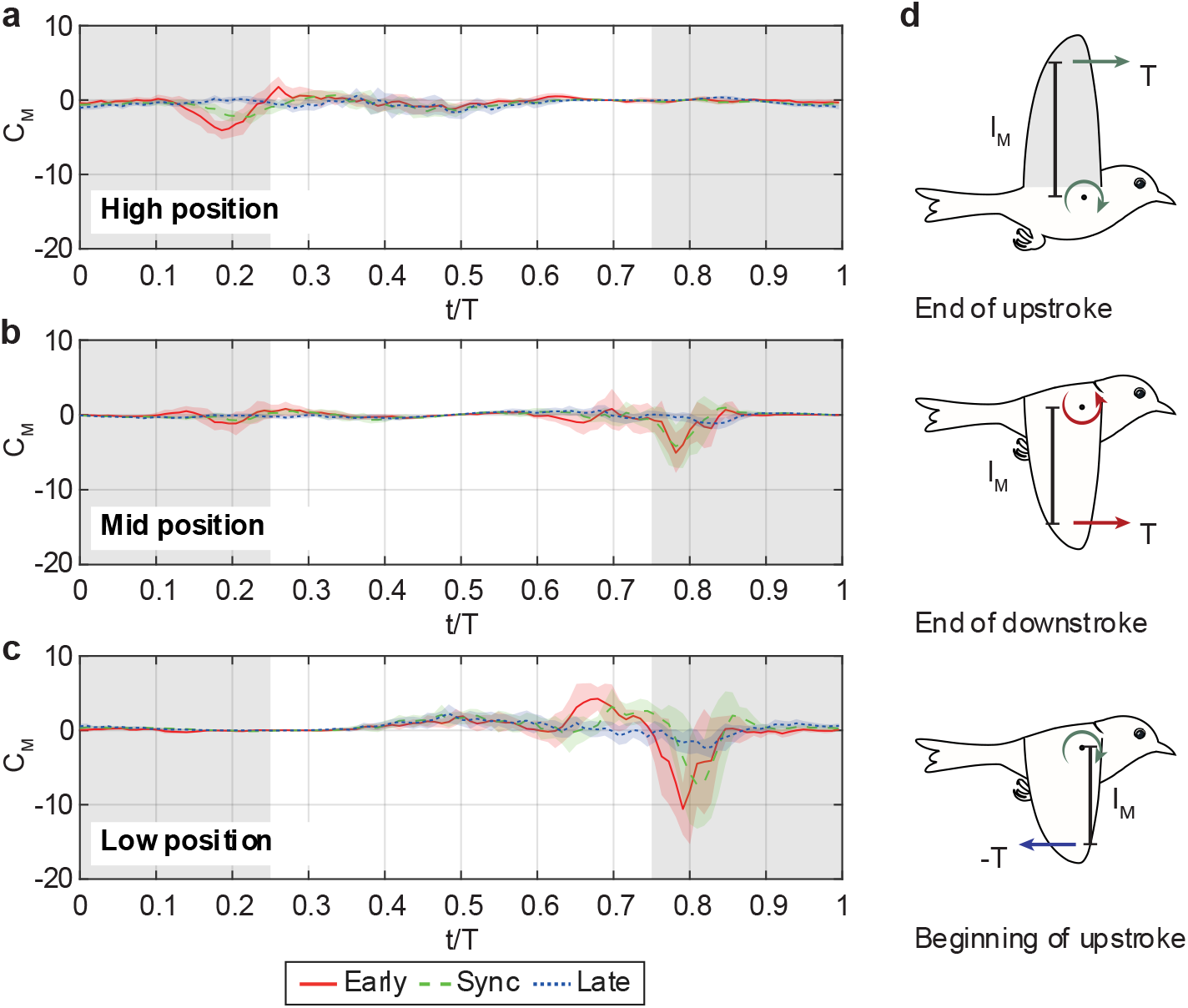
Pitch torque (*C*_M_) acting on the body varied with kinematics. (a) When flapping the wing in a high position, the thrust (*T*) generated during the upper turning point created a pitch down torque, while thrust generated at the lower turning point had little influence on *C*_M_. (b) When the wing was in the mid position, *T* during both turning points resulted in increased *C*_M_ magnitude. (c) With the wing in the lower position, the lower turning point had a large impact on *C*_M_, both during the end of the downstroke and the beginning of the upstroke. (d) Illustration of the moment arm (*l*_M_) and *T* creating the pitch torque on the body during the turning phases of the wingbeat.

## Discussion

Using a robotic wing to control individual aspects of wing kinematics during forward flight conditions, we have shown that both mean wing elevation (*MWE*) and the timing of wing pitch (*TP*) at up- and downstroke transitions influence the magnitude, direction and efficiency of aerodynamic force generation. Importantly, these effects are not independent, and the same *TP* shift can either enhance or reduce lift depending on *MWE*, demonstrating that *TP* must be interpreted in the geometric and aerodynamic context set by wing elevation. This parameter interaction was strongest for lift, force direction, and the efficiency measures. Depending on whether the goal is thrust production, weight support, or energetic efficiency, different regions of the *MWE*–*TP* control space should be selected. This reveals a rarely considered complexity of flight control that is likely available to flying animals and offers novel coupled control space for designing flapping drones. During flapping flight, the distance and angle between the wings vary throughout a wingbeat. When the wings are close, either above or below the body, they interact aerodynamically (52). The wings have mirror pressure distributions (52,53), which affect their aerodynamics. When the wings move apart at the start of the downstroke, this results in a lower pressure between them than would be found if they operated on their own, which tends to increase lift (54,55). In our measurements, we find the expected increase in *C*_L_ when the wings are high, but the extra lift mainly comes at the end of the downstroke rather than at the start (Fig. 2a). We interpret this as a consequence of the downstroke ending in a position where the wing is close to the horizontal plane and pitch up during this phase increases the angle of attack and the lift production. In addition, when the wings are high and the aerodynamic interaction between the wings is expected to be significant, the wing elevation results in the force generated from each wing being directed sideways, contributing relatively little to vertical force. When *MWE* instead is low and the wings come together at the end of the downstroke, we observed an increase in thrust, which we interpret as resulting from an increased pressure between the wings. This is similar to when wings are in ground effect (42,43,46). Earlier studies on flapping flight in ground effect have suggested that how the wings are pitched when they interact with the ground affects the performance, which is thought to explain higher than predicted energy savings for bats (46) and modelled birds (56). In agreement with these studies, we found that the amount of lift or thrust at the end of the downstroke depends on the pitch angle of the wings when they approach each other. If the leading edges of the wings meet first, as when using a delayed pitch, this results in more thrust than during the synchronized case when the wings have no pitch at the turning point. We suggest this to be the result of the high pressure on the bottom wing surfaces accelerating air backwards during the wing-wing interaction. If instead the trailing edges meet first, as when pitching up early, the high –pressure on the bottom wing surface would accelerate air forwards, which would then generate negative thrust during this phase (Fig. 2). Our results thus support the notion that the attitude of the wings, or their pitch angle, during the wing-wing interaction makes a significant impact on the outcome of the interaction. This also proposes the opportunity that adjusting *TP* differently for the upper and lower transitions phases will add further controllability.

An interesting finding is that both the *PF* and *COT* suggest that efficient force generation, in our setup, happens when *TP* occurs late and *MWE* is in the range between the mid and high position. Looking at the components of the efficiency measures we see that this result is largely driven by a low *CP* for later *TP* and relatively high force generation (*C*_L_ and *C*_T_). Consequently, unless necessary for force balance or maneuver/stability reasons, we would suggest animals to favor flapping their wings relatively symmetrically around the horizontal plane. Overall, our findings suggest that within our parameter space, *C*_L_ is maximized by flapping the wings elevated and with an early *TP*, while for maximizing *C*_T_ the wing should be in a mid-to-low position with a delayed *TP*. If the goal is to change the direction of the force generated (i.e. *T*/*L*), a high wing position with an early *TP* tilts the force upwards the most and a low wing position with a synced *TP* tilts the force forwards the most (Fig. 5). This means that simple, but relatively understudied, kinematic changes can be used to control flight. One of the key factors with the parameters we have studied here is that they, in addition to being allowed to vary between wingbeats, can be adjusted within a single wingbeat (Fig. 2). For birds and bats, these results suggest that *TP* and *MWE* may function as rapid within-wingbeat control variables, allowing animals to redistribute force between weight support, propulsion, and body-pitch (see below) regulation without large changes in gross wing trajectory. A prediction based on these findings is that we will be likely to find large variation in how animals generate maneuvers or stabilize flight, depending on the phase of the wingbeat they decide to adjust their trajectory. This will at least apply to animals with relatively low wingbeat frequency compared to the speed of their control system (sensors and processing), such as birds and bats, since they would benefit relatively more from course corrections within wingbeats. Indeed, one of the few studies looking at the aerodynamics of maneuvers actually used by flying vertebrates (bats) have showed a variety of mechanisms (including varying the *MWE* of the left and right wing separately) depending on the phase of the wingbeat when the maneuver is initiated (10).

**Figure 5.**
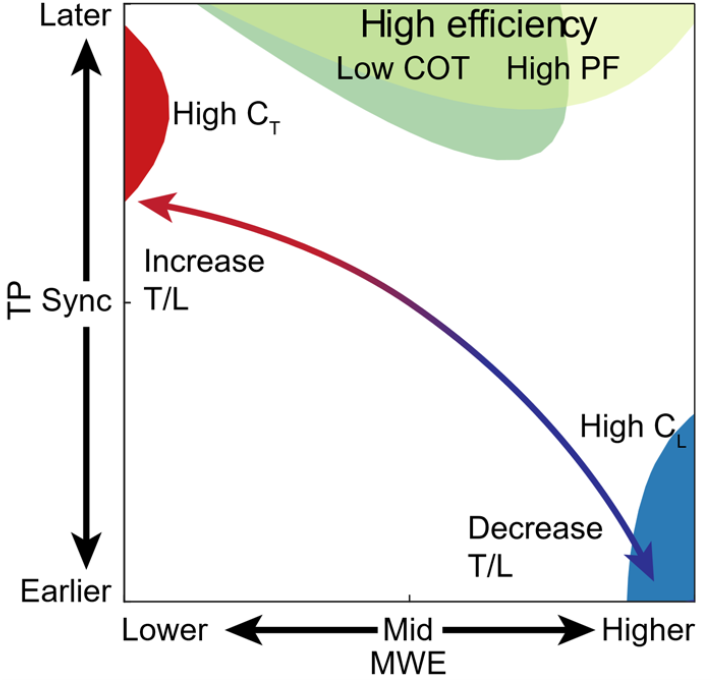
Kinematic control map summarizing tradeoffs across the sampled *TP*–*MWE* parameter space. High *MWE* with early *TP* favors lift and an upward-tilted resultant force, whereas low *MWE* with late *TP* favors thrust and a more forward-directed force. Energetic performance is best near mid-to-high *MWE* with late *TP*. The map emphasizes that *TP* and *MWE* define a control space in which different aerodynamic objectives are favored in different regions.

Our results were obtained for a flapping robot fixed to the wind tunnel. As a consequence, our robot generated net thrust and variation in lift, while an equilibrium with zero net thrust (thrust and drag cancel each other) and lift equaling weight is quickly reached in free flying animals and drones. Interpreting the robot’s force generation in a free flight context, we infer that the high thrust coefficients would lead to acceleration and an increase of drag before stabilizing at a higher flight speed. Conversely, high values of the lift coefficients suggest enabling flight at lower speeds, while low values require higher speeds to obtain weight support. Our results thus suggest that low *MWE* and late *TP* may provide suitable kinematics for fast flight, where low values of the lift coefficient can produce sufficient weight support, and more thrust is required because of the increased profile and parasitic drag. On the other hand, we suggest that high *MWE* combined with early *TP* should be preferred at slow flight, where weight support requires high lift coefficients, and lower speed produce less drag/thrust requirements. Interestingly, this proposed pattern with high *MWE* at low speed and low *MWE* at high speed has been demonstrated in some freely flying birds(6,7) and bats(4,8,38) suggesting animals have already adapted this strategy. By contrast, data on how animals adjust *TP* in forward flight is largely lacking, while our results identify *TP* as a potentially important control variable.

Another flight control mechanism, which is linked to the transition phases, involves the relatively large variation in thrust when we change the *TP* (Fig. 2). Modulating thrust production during the phases when the wings reach their most elevated or most depressed positions provides an effective means of controlling body pitch (Fig. 4). Changes in the *MWE* simultaneously alter both the magnitude of thrust (Figs 1 and 2), through modifications to wing–wing aerodynamic interactions, and the moment arm of this force, by shifting the vertical position of the wings at the phase when thrust is produced. Together, these effects determine the pitch torque acting on the body. This mechanism thus provides a tailless pitch control that can be used to stabilize flight when flying without a tail (37), but also to initiate maneuvers (57). More generally, this implies that it is possible to generate yaw torques by employing different *TP* strategies for the left and right wings, while left–right asymmetries in *MWE* can be used to produce roll torques (10,58). These control strategies can potentially be applied in concert with other mechanism, including active wing morphing (59,60), resulting in a flexible control mechanism that can be applied within a single wingbeat and with a short response time. Understanding how different species utilize these mechanisms to achieve maneuvers, depending on wingbeat phase and strength of the maneuver, represents an exciting avenue for future research.

## Conclusion

Our results demonstrate that previously overlooked kinematic strategies can substantially influence the direction, magnitude, and efficiency of aerodynamic forces generated by flapping wings during forward flight. Both mean wing elevation and the timing of wing pitch independently affect aerodynamic performance, but notably, we also uncover a strong interaction effect between them. This suggests that these two kinematic parameters can be combined to enhance force production or improve efficiency further than either parameter alone. Physically, we suggest this interaction relates to how *TP* and *MWE* alter how the wings present their pressure surfaces during close wing–wing interactions. Moreover, we show that these kinematic adjustments determine when thrust is produced within the wingbeat, thereby modulating the pitch torque acting on the body. To improve pitch torque authority, our results suggest that transition phases with large moment arm should be used. This provides a tailless mechanism that animals could exploit to stabilize flight or initiate maneuvers. By treating *TP* and *MWE* as coupled kinematic control variables, this study shows that small within-wingbeat adjustments can control force magnitude, direction, efficiency, and pitch in different ways. The results therefore provide both a mechanistic framework for interpreting animal wing motions and a control-oriented design map for bio-inspired flapping systems.

## Acknowledgement

The research was funded by grants from the Crafoord foundation (20170584) and the Swedish Research Council (2017-03890 and 2022-02850 to L.C.J.). The PIV equipment was financed by an infrastructure grant from Lund University (https://www.lu.se/) to Anders Hedenström, Per Henningsson and L.C.J..

## Notes

### Competing Interest Statement

The authors have declared no competing interest.

### Summary of Updates

Added figures to the manuscript and supplemental information and revised the text. Figures have been updated and additional statistical analysis has been performed.

