## Supplemental Figures for "Wing pitch timing and wing elevation modulate forces and body pitch in forward flapping flight"

### Supplemental information

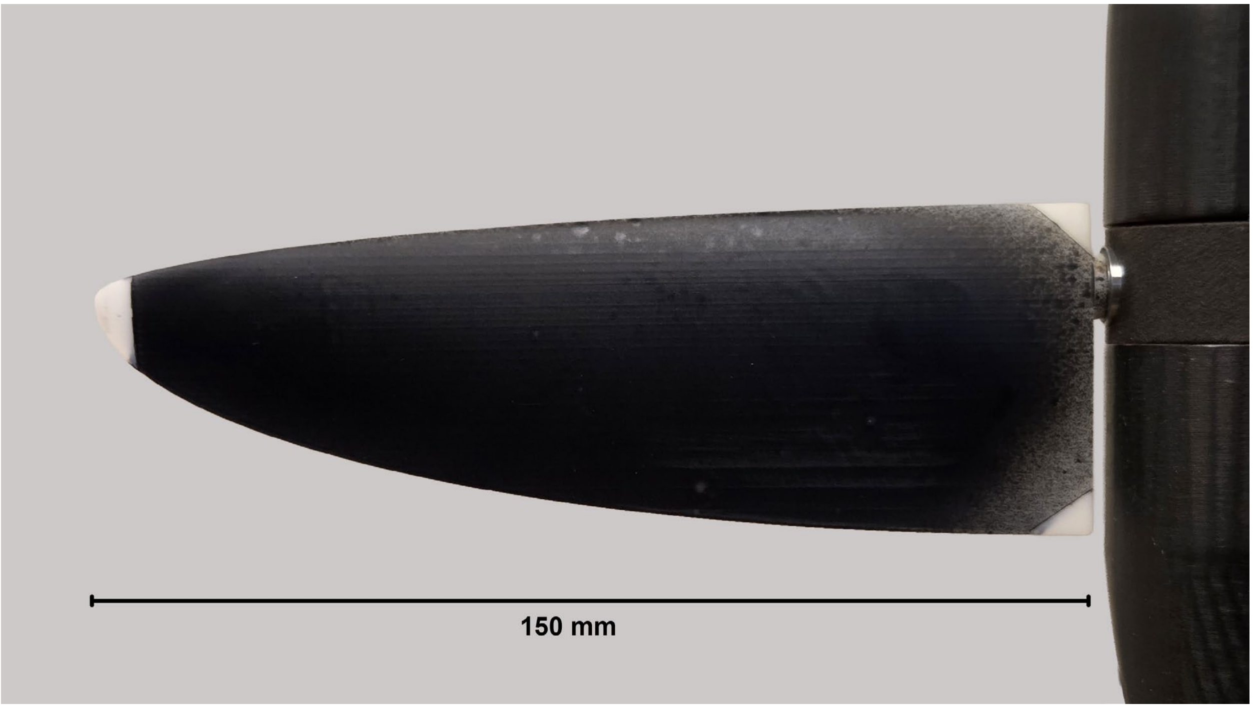


**Figure S1.** Wing used on the robot.

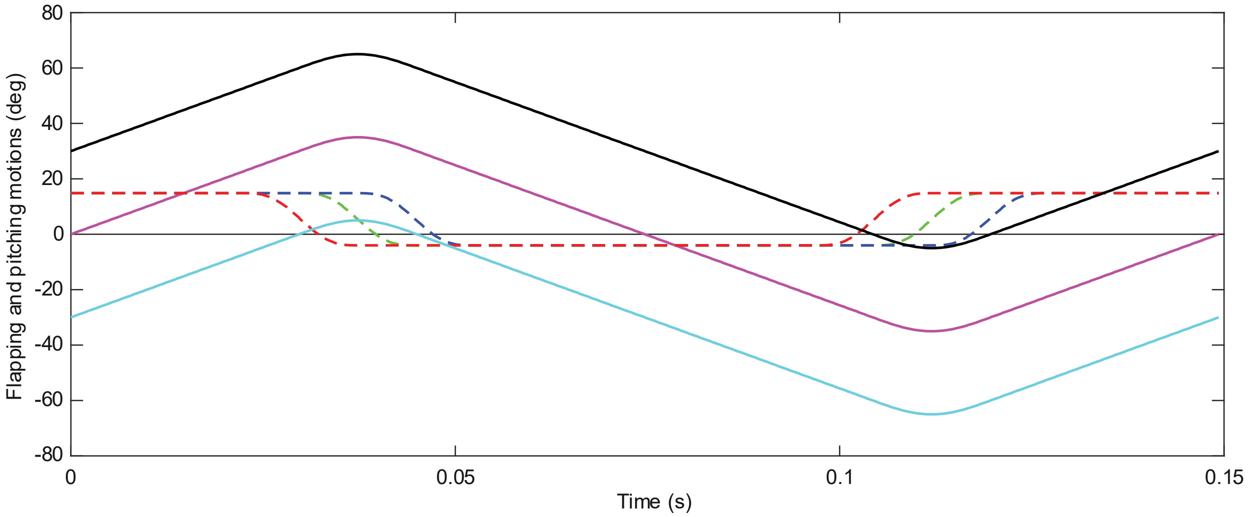


**Figure S2.** Temporal variation in flapping angle (black-high, magenta-mid and cyan-low) and pitching angle (red-early, green-synchronized and blue-late) over time.

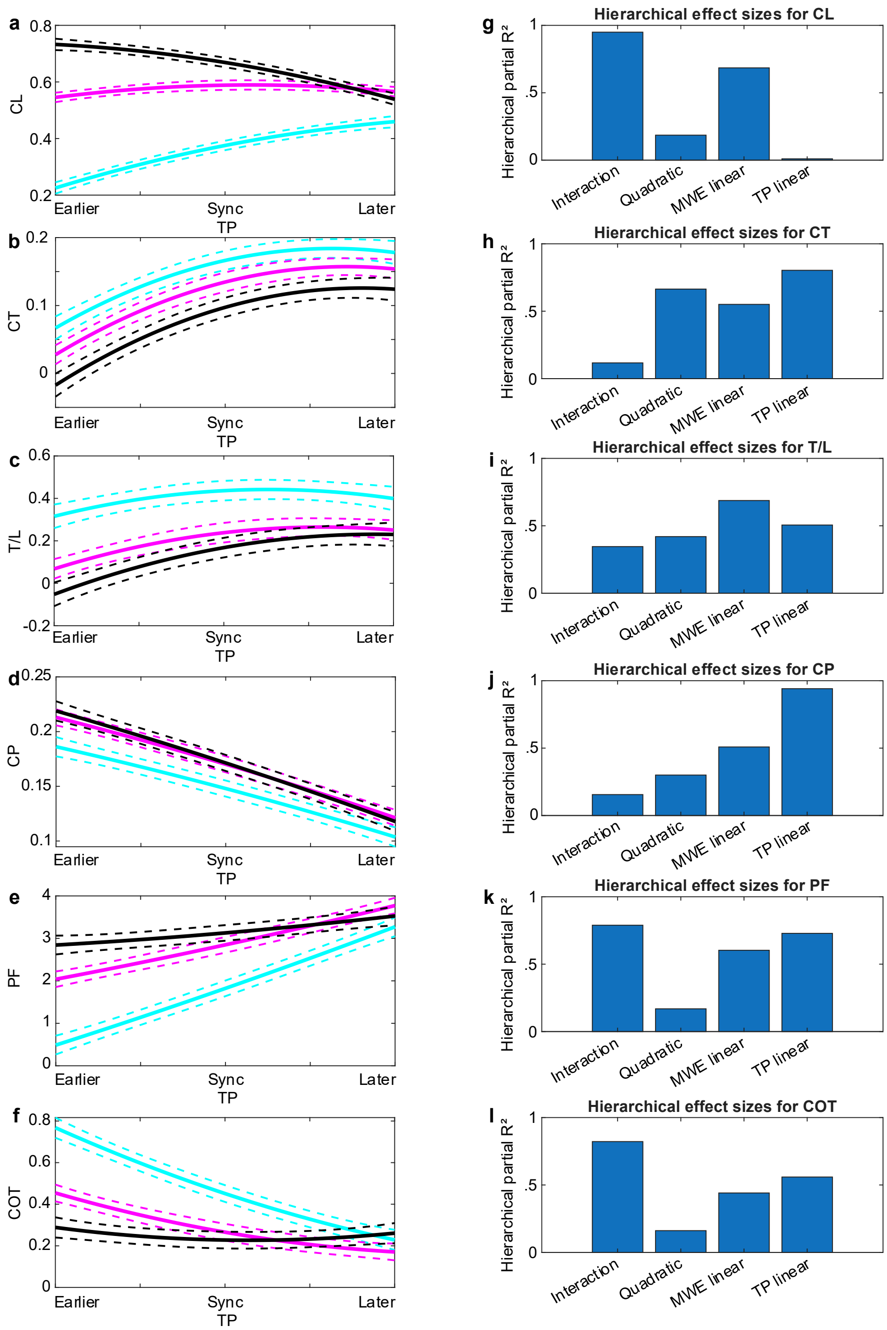


**Figure S3**. Interaction plots of the fitted quadratic GLM (a-f) visualize whether the effect of pitch timing remained constant across mean wing elevation or instead changed in magnitude or direction. Parallel curves indicate weak interaction, whereas non-parallel or crossing curves indicate that the aerodynamic effect of *TP* depends on *MWE*. Colors represent *MWE* with black = high, magenta = mid and cyan = low. (g-l) To compare the relative contributions of linear, nonlinear, and interaction structure, we performed hierarchical nested-model comparisons and quantified the incremental explanatory power of each block of terms.

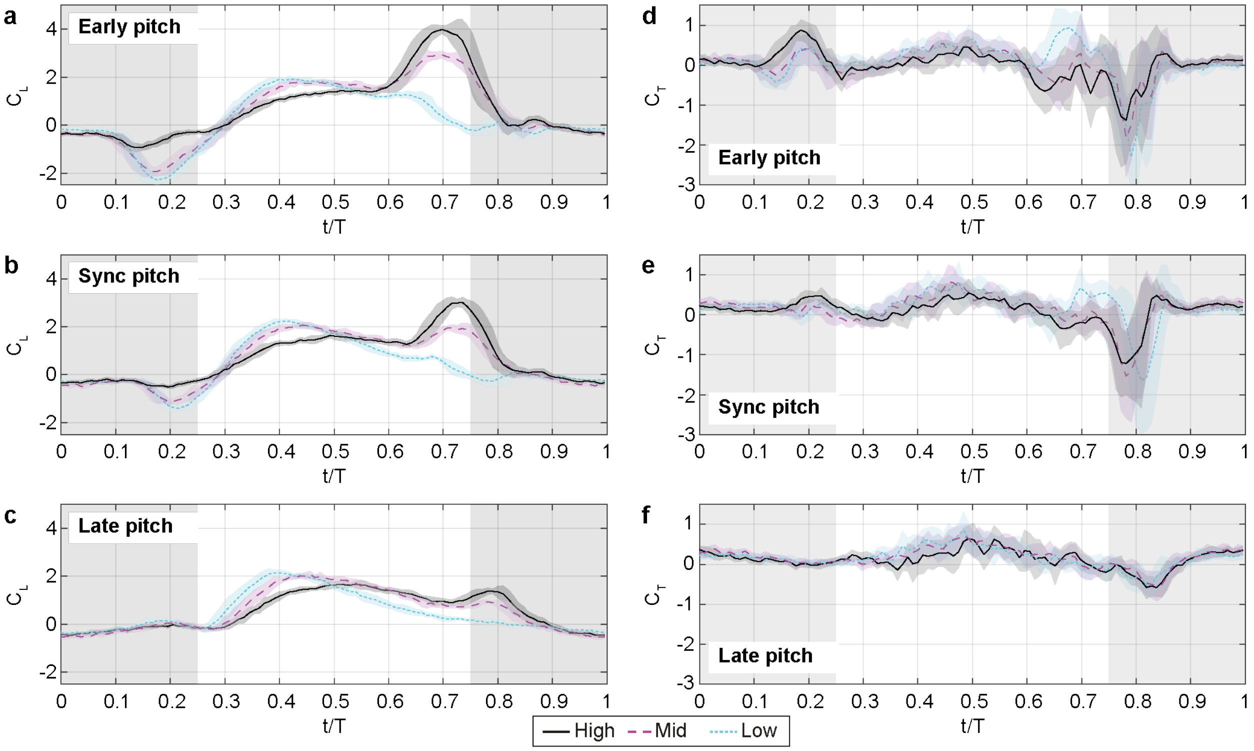


**Figure S4.** Instantaneous force coefficients through the flapping cycle. The left panels (**a-c**) show the impact on $C_L$ and the right panels (**d-f**) show the impact on $C_T$ for the different wing positions (*MWE* = High, Mid and Low) for each of the timing of the pitch (*TP* = Early, Sync and Late). In each curve, the line represents the phase-averaged value over the three repeats and four wingbeat of each case, and the shaded area has a width of two standard deviations centered around the mean. The data is the same as in Fig. 2 but divided in the alternate way to facilitate interpretation. Shaded background indicates the upstroke.

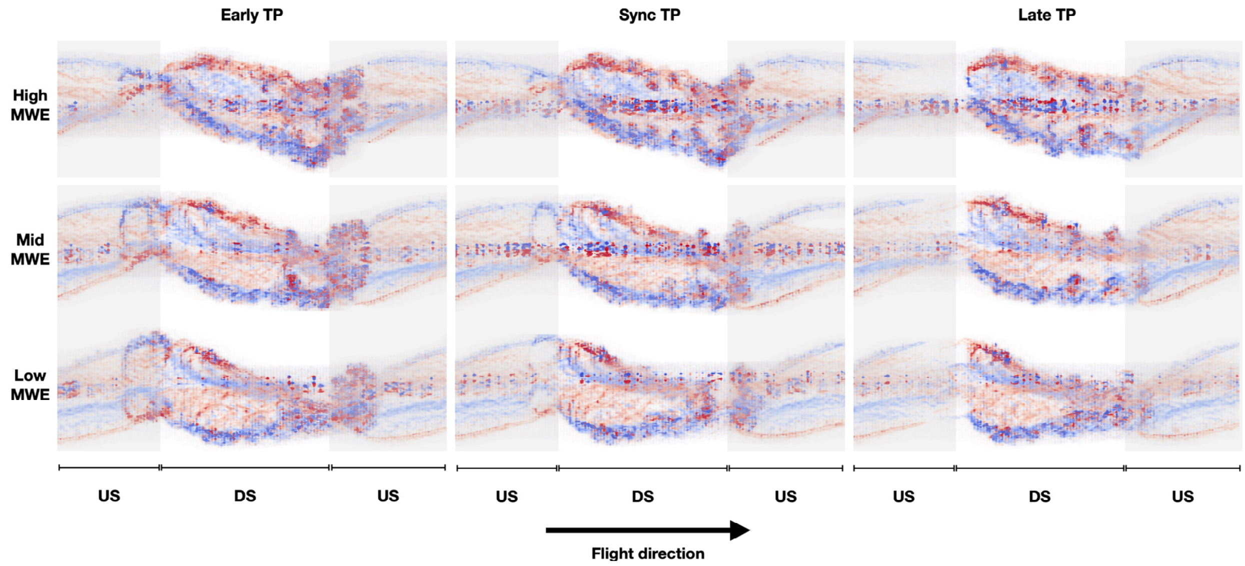


**Figure S5.** Volume rendering of the reconstructed vorticity field for one of each tested case. The magnitude of the vorticity vector is used for the rendering, and it is colored towards red or blue depending on the sign of the streamwise component of the vorticity. The upstrokes are highlighted with shaded areas.
